# Using unlabeled information of embryo siblings from the same cohort cycle to enhance in vitro fertilization implantation prediction

**DOI:** 10.1101/2022.11.07.515389

**Authors:** Noam Tzukerman, Oded Rotem, Maya Tsarfati Shapiro, Ron Maor, Marcos Meseguer, Daniella Gilboa, Daniel S. Seidman, Assaf Zaritsky

## Abstract

High content time-lapse embryo imaging assessed by machine learning is revolutionizing the field of in vitro fertilization (IVF). However, the vast majority of IVF embryos are not transferred to the uterus, and these masses of embryos with unknown implantation outcomes are ignored in current efforts that aim to predict implantation. Here, we explore whether, and to what extent the information encoded within “sibling” embryos from the same IVF cohort contribute to the performance of machine learning-based implantation prediction. First, we show that the implantation outcome is correlated with attributes derived from the cohort siblings. Second, we demonstrate that this unlabeled data boosts implantation prediction performance. Third, we characterize the cohort properties driving embryo prediction, especially those that rescued erroneous predictions. Our results suggest that predictive models for embryo implantation can benefit from the overlooked, widely available unlabeled data of sibling embryos by reducing inherent noise of the individual transferred embryo.

**Significance statement:** We use in vitro fertilization (IVF) as a model to study the effect of genotypic and environmental variation on phenotype and demonstrate a potential translational application. This is achieved by associating the implantation potential of transferred embryos and the visual information encoded within their non-transferred “sibling” embryos from the same IVF cohort, and establishing that these cohort features contribute to consistent improvement in machine learning implantation prediction regardless of the embryo-focused model. Our results suggest a general concept where the uncertainty in the implantation potential for the transferred embryo can be reduced by information encapsulated in the correlated cohort embryos. Since the siblings’ data are routinely collected, incorporating cohort features in AI-driven embryo implantation prediction can have direct translational implications.

## Introduction

Phenotypic variation is inherent in every biological system. A phenotype is determined by a combination of genetic and environmental factors. For example, the proportion of shared genetic background between human siblings can explain most of the variability in height [1], while environmental factors can affect gut microbiota composition [2], gene expression and disease susceptibility [3], and even lead to phenotypic variation in monozygotic (aka “identical”) twins [4]. Thus, confinement of the genetic and environmental variability, for example, by considering siblings raised under similar conditions, can lead to reduced phenotypic variability, i.e., increased phenotypic similarity. We hypothesized that such phenotypic correlations can provide predictive value regarding an individual’s future phenotypic state by considering the phenotypic states of its siblings.

Specifically, during in vitro fertilization (IVF) a cohort of “sibling” oocytes, all sharing the same “parents’’ within the same IVF treatment, are fertilized and incubated for up to six days under the same laboratory conditions before one or a few embryos from the cohort are selected for transfer into the uterus. IVF embryo phenotypes are heavily affected by genetic [5] and environmental [6] factors, and thus the genetic and environmental variability is minimized for siblings from the same cohort.

Recent advances in time-lapse video microscopy for live embryo imaging has transformed IVF into a data-intensive field. This has led to innovative attempts to automatically and unbiasedly estimate the implantation potential of embryos based on algorithmic assessment of their visual phenotypes and/or developmental trajectory [7,8,9,10,11,12]. Specifically, supervised machine learning has emerged as a powerful approach where features, computationally extracted from embryo images with known implantation outcomes, are used to train computational models to predict implantation [13,14,15,16]. These models reach performance comparable and even exceeding those of embryologists [17,18,19].

However, from the available cohort, only one or at most two embryos are selected for transfer to the uterus. This poses a major limitation for machine learning-based embryo selection approaches because the implantation potential of the majority of deselected embryos remains unknown [16]. Thus, the number of embryos with known implantation outcomes available for training is severely limited. Recent studies have attempted to overcome this limitation by including “unlabeled” embryos in their model training schemes, specifically focusing on the subgroup of embryos found unsuitable for transfer to the uterus due to their poor morphological appearance and presumed limited implantation potential [2021]. Other studies used the morphological annotations of embryos found unsuitable for transfer to the uterus to train models to predict the value of these morphological measurements as a readout for successful implantation potential [22,23]. However, none of these studies fully capitalized on and systematically assessed the potential of using the association between “sibling” embryos from the same cohort to enhance implantation prediction accuracy.

We asked whether we can take advantage of cohort-derived information to train a machine learning model tasked with predicting implantation? We hypothesized that cohort embryos share information relevant for implantation prediction. Thus, using the full extent of the available unlabeled data in the cohort may provide further statistical discriminative power. Indeed, a few earlier studies provide evidence supporting the notion that cohort siblings encapsulate information that correlate with the transferred embryo’s quality and implantation outcome, such as cohort size [24,25,26,27], sibling blastocyst development [2829], or a combination of cohort-specific variables [26]. Here, we explicitly assess the contribution of sibling information to embryo implantation prediction by systematically evaluating different models trained with or without information from the cohort. We demonstrate that the unlabeled cohort embryos contribute to the prediction. Our results imply that artificial intelligence (AI)-based embryo assessment can benefit from the widely available, and currently ignored, correlated data in the cohort’s siblings. We characterize the specific properties of the cohort that contribute to the prediction and show that different cohort features can be exploited to enhance performance of different models in varying contexts. Altogether, we suggest that considering the correlated properties of sibling embryos aid in canceling the inherently noisy single embryo prediction.

## Results

### Embryos from the same cohort are phenotypically correlated

Our data included information derived from 2089 transferred embryos collected from 1605 IVF cycles that included 1176 implanted blastocysts (*positive embryos*), 913 non-implanted blastocysts (*negative embryos*), and 14105 sibling embryos that were not transferred (Methods, Table S1). The timing of 7 hallmark stages in embryo development, termed morphokinetic events, were manually annotated based on time-lapse observation of the developing embryos (Fig. 1A, Table S1, Methods). These morphokinetic events are considered key in proper embryo development and were shown to be correlated with implantation potential [30,10]. The timing of morphokinetic events from fertilization and the time intervals between consecutive morphokinetic events were more similar among sibling embryos than for randomly selected non-sibling embryos, indicating lower intra-cohort variability (Fig. S1A-F - timing of morphokinetic events, Fig. 1B-F - time intervals between consecutive morphokinetic events, Fig. 1G - normalized distances between multivariate representations of all time intervals between consecutive morphokinetic events, Fig. S1G - normalized distances between multivariate representations of all morphokinetic events). To verify that this higher intra-cohort (i.e., between siblings) similarity is not an artifact due to the random selection of embryos, we compared intra- versus inter-cohort similarity in the morphokinetic profiles of embryos with similar morphological qualities. We considered the manually annotated Gardner and Schoolcraft alphanumeric quality scoring scheme as a proxy for embryo morphological-based quality, which is based on the assessment of three parameters: blastocyst expansion status, morphology of the inner cell mass (ICM), and morphology of the trophectoderm (TE) [31,32] (Fig. 1H). We considered all possible ordered combinations of triplets of embryos that include a pair of sibling embryos from the same cohort and a third non-sibling embryo from a different cohort, where all three embryos had the same Gardner scores (Fig. 1I, Methods). This analysis established that morphokinetic multivariate profiles of sibling embryos are more similar than non-sibling embryos and that this similarity is not a mere consequence of a morphologic similarity between siblings at the blastocyst stage (Fig. 1J). Altogether, this data supports the notion that sibling embryos share similar phenotypic properties.

**Figure 1.**
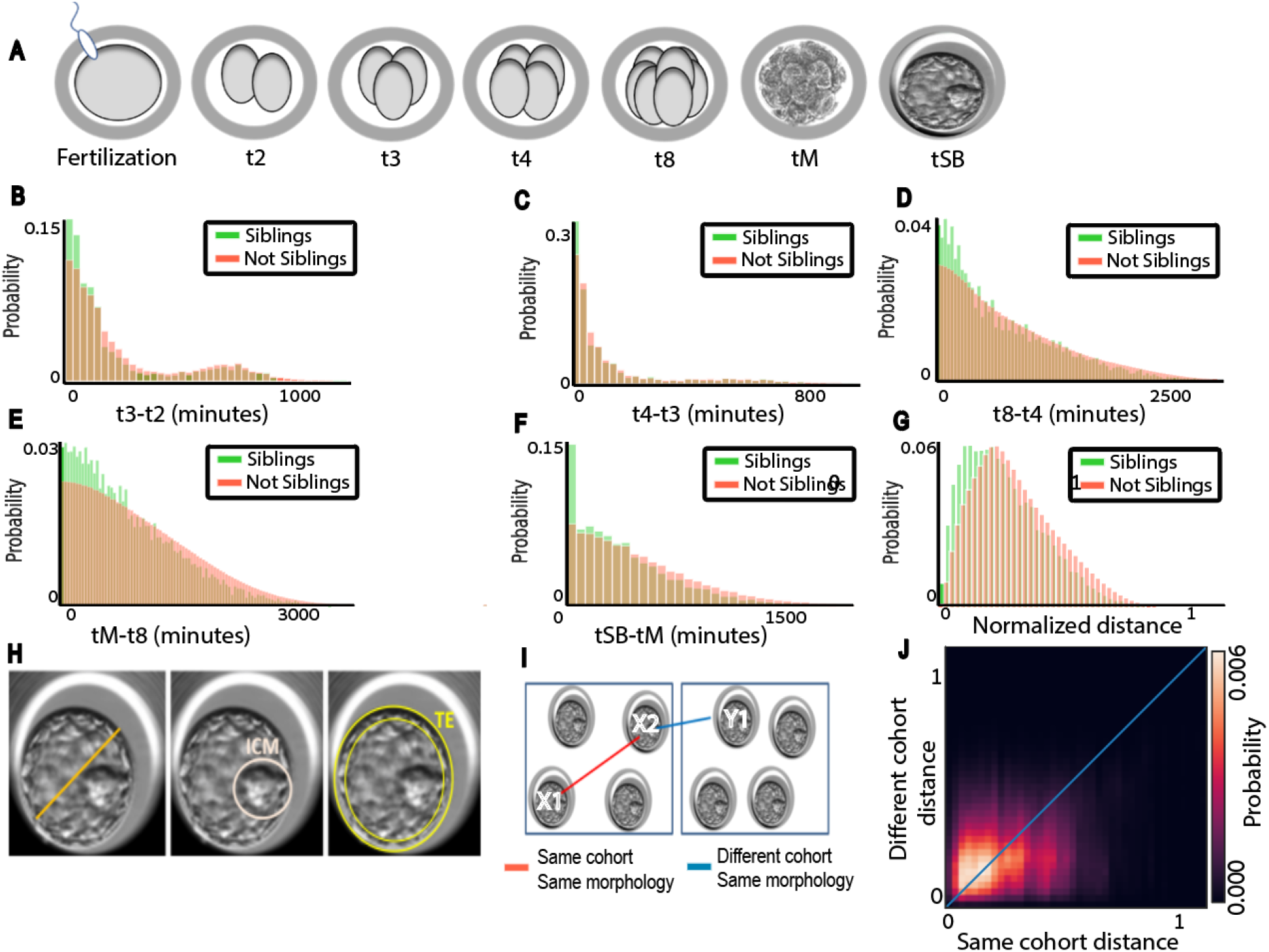
Sibling embryos from the same cohort are more similar than non-siblings in terms of their morphokinetic properties. (**A**) Schematic sketch. Morphokinetic features: cell division to the 2, 3, 4 and 8-cell stage (t2, t3, t4, t8), the compaction of the morula - a day-3 development stage (tM) and the start of blastulation (tSB) - a day-5 development stage. (**B-G**) Distribution of the difference in time intervals, in minutes (B-F) or normalized distance (G), between consecutive morphokinetic events compared across siblings versus not siblings embryo pairs. N embryos = 16194. N cohorts = 1605. N positive cohorts = 928, N negative cohorts = 677. (**B**) Mean (standard deviation) of distances between t3-t2 intervals was 174.03 (214.7) for sibling embryos versus 202.26 (218.7) for not-sibling embryos, Mann-Whitney-U signed rank test p-value < 0.0001. (**C**) Mean (standard deviation) of distances between t4-t3 intervals was 146.48 (220.56) for sibling embryos versus 158.75 (225.52) for not-sibling embryos, Mann-Whitney-U signed rank test p-value < 0.0001. (**D**) Mean (standard deviation) of distances between t8-t4 intervals was 594.58 (531.66) for sibling embryos versus 684.09 (569.68) for not-sibling embryos, Mann-Whitney-U signed rank test p-value < 0.0001. (**E**) Mean (standard deviation) of distances between tM-t8 intervals was 691.03 (546.57) for sibling embryos versus 815.16 (602.9) for not-sibling embryos, Mann-Whitney-U signed rank test p-value < 0.0001. (**F**) Mean (standard deviation) of distances between tSB-tM intervals was 344.50 (336.89) for sibling embryos versus 434.79 (372.23) for not-sibling embryos, Mann-Whitney-U signed rank test p-value < 0.0001. (**G**) Mean (standard deviation) of normalized distances between all morphokinetic features time intervals was 0.241 (0.14) for sibling embryos versus 0.28 (0.15) for not-sibling embryos, Mann-Whitney-U signed rank test p-value < 0.0001. (**H**) Predefined criteria of blastocyst quality according to the Gardner three-part scoring scheme. From left to right: Blastocyst expansion – volume and degree of expansion of the blastocyst cavity (ranked 1-6). Morphology of the Inner cell mass (ICM) –size and compaction of the mass of cells that eventually form the fetus (ranked A-D). Morphology of the trophectoderm (TE) – number and cohesiveness of the single cell layer on the outer edge of the blastocyst that eventually forms the placenta (ranked A-D). (**I**) Schematic sketch of the analysis comparing embryo triplets: two from the same cohort (annotated X1 and X2), and two from a different cohort (X2 and Y1), where all embryos have the same Gardner annotations (similar morphological quality). (**J**) Mean (standard deviation) of normalized distances between all morphokinetic features time intervals was 0.224 (0.141) for sibling embryos with similar Gardner scores versus 0.263 (0.146) for not-sibling embryos with similar Gardner scores, Mann-Whitney-U signed rank test p-value < 0.0001. N = 1,654,732 ordered triplets.

### Cohort properties correlate with implantation outcome

The phenotypic similarity between siblings from the same cohort raises the hypothesis that morphological and morphokinetic properties of sibling embryos are correlated to the implantation outcome. To test this hypothesis we compared the distribution of several cohort-related properties for cohorts that included successfully implanted embryos (*positive cohorts*) versus those cohorts where the transferred embryo/s failed to implant (*negative cohorts*). First, we validated that positive cohorts contained more embryos than negative cohorts (Fig. 2A), and that the fraction of sibling embryos within a cohort (not including the transferred embryo/s) reaching blastulation was larger in positive cohorts (Fig. 2B). These results are in agreement with previous reports for cohort size [25,27,33,24,26] and for siblings blastocyst development [28,29]. Each of the three Gardner morphological scores was elevated for sibling embryos, in positive cohorts compared to negative cohorts (Fig. 2C-E). While the enrichments of higher quality cohort properties in positive cohorts were relatively small for each cohort property, they were all consistent toward favoring higher quality cohort properties in positive cohorts. Cumulatively, these results conclude that morphological properties of “sibling” embryos within a cohort, that were not transferred, are associated with the implantation potential of the embryo that was transferred within that cohort. These results establish that properties derived from sibling embryos correlate with the clinical outcome of their sibling transferred embryo.

**Figure 2:**
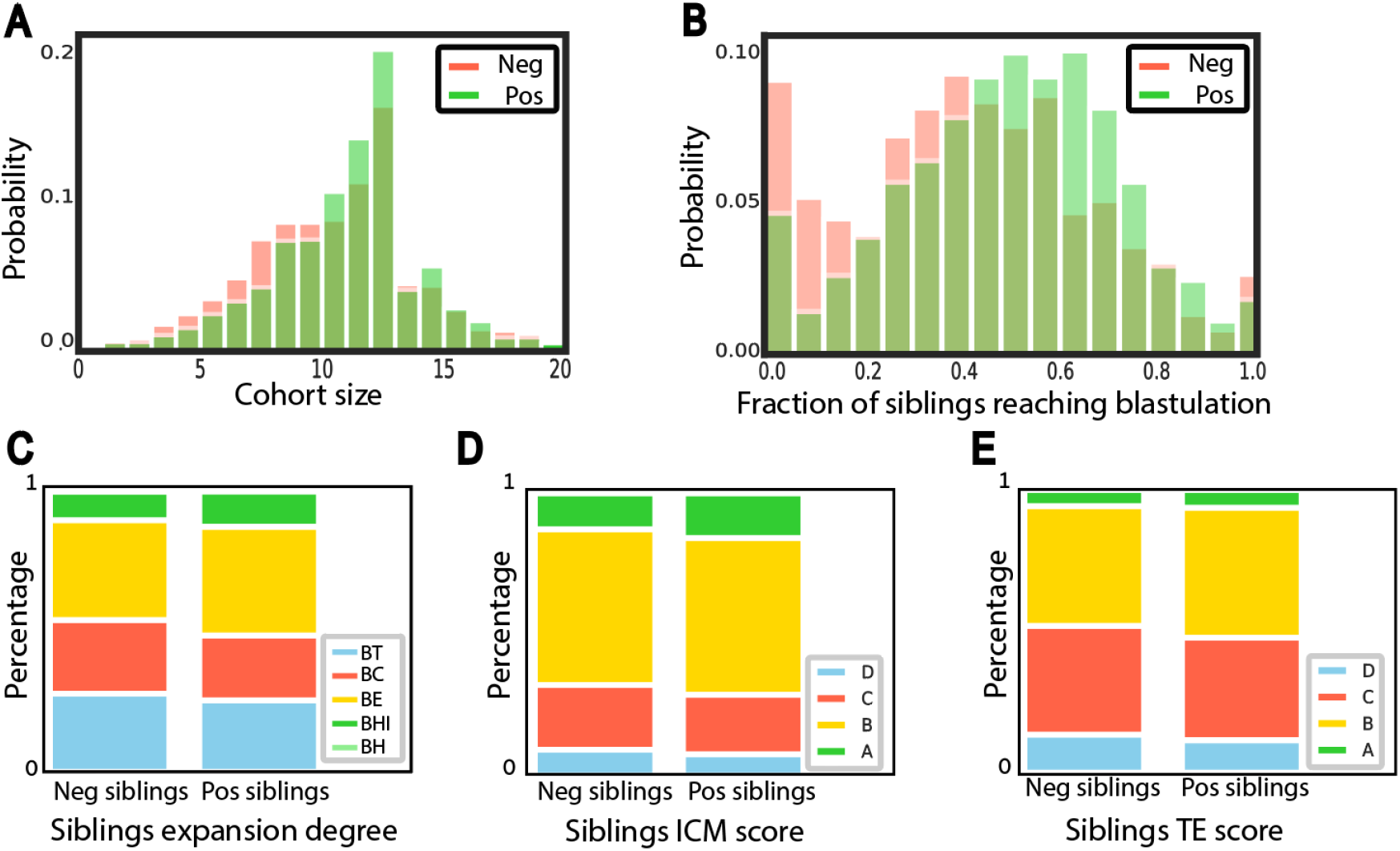
Siblings in positive cohorts are of higher morphological quality than those in negative cohorts. (**A-B**) Distribution of the cohort size (i.e., number of sibling embryos in a cohort) (A) or the fraction of embryos within a cohort (not including the transferred embryo/s) to develop into a blastocyst (B) compared across positive versus negative cohorts. N transferred blastocysts =2089. N implanted blastocyst = 1176, N non-implanted blastocyst = 913. N cohorts = 1605. N positive cohorts = 928, N negative cohorts = 677. (**A**) Mean (standard deviation) cohort size was 10.56 (3.19) for positive cohorts versus 9.9 (3.36) for negative cohorts, Mann-Whitney-U signed rank test p-value < 0.0001. (**B**) Mean (standard deviation) fraction of sibling embryos within a cohort (not including the transferred embryo/s) reaching blastulation was 0.49 (0.22) for positive cohorts versus 0.41 (0.25) for negative cohorts, Mann-Whitney-U signed rank test p-value < 0.0001. (**C-E**) Distribution of manually annotated Gardner scores of cohort embryos (not including the transferred embryo/s) across positive versus negative cohorts. N transferred blastocysts=1936 embryos. N implanted blastocysts = 1141, N non-implanted blastocysts = 795. (**C**) Expansion degree: BT-early blastocyst, BC - full blastocyst, BE-expanded blastocyst, Bhi - hatching blastocyst, BH - hatched blastocyst. The total number of BH embryos in our dataset is 3 for implanted blastocysts and 3 for non-implanted blastocysts, and thus cannot be seen in the graph. Corresponding Mann-Whitney-U signed rank tests on the null hypothesis that the two schemes were drawn from the same matched distribution p-value < 0.0001. (**D**) Inner cell mass (ICM): ranked from A (high quality) to D (low quality). Corresponding Mann-Whitney-U signed rank tests on the null hypothesis that the two schemes were drawn from the same matched distribution p-value < 0.0001. (**E**) Trophectoderm (TE): A (high quality) to D (low quality). Corresponding Mann-Whitney-U signed rank tests on the null hypothesis that the two schemes were drawn from the same matched distribution p-value < 0.0001.

### Cohort properties contribute to implantation prediction

Given that cohort properties were correlated to the implantation outcome, we hypothesized that inclusion of cohort-derived features can enhance the prediction power of a machine learning model initially trained without cohort information (Fig. 3A). To assess the generality of this idea we trained several distinct machine learning models for the prediction of implantation outcome, each of these models was trained without and with cohort-derived features. The performance of each pair of models, without or with cohort features, was compared to assess the contribution of the cohort information. The first model was morphology-based, and trained on manually annotated Gardner scores (Methods). The second model was morphokinetics-based [7], and trained on manually annotated key morphokinetic events (Methods). The third model combined morphology, morphokinetics and the oocyte age, where the latter is widely accepted as correlated with implantation success [3435] (Methods). Seventeen cohort-derived features were calculated from the siblings of the transferred embryo at test. These included cohort size, fraction of sibling embryos reaching blastulation, and features encoding the siblings’ Gardner scores (Methods). The performance of each of these three models was improved by incorporating cohort-derived features, as measured by receiver operating characteristic (ROC) area under the curve (AUC) (Fig. 3B-D).

**Figure 3:**
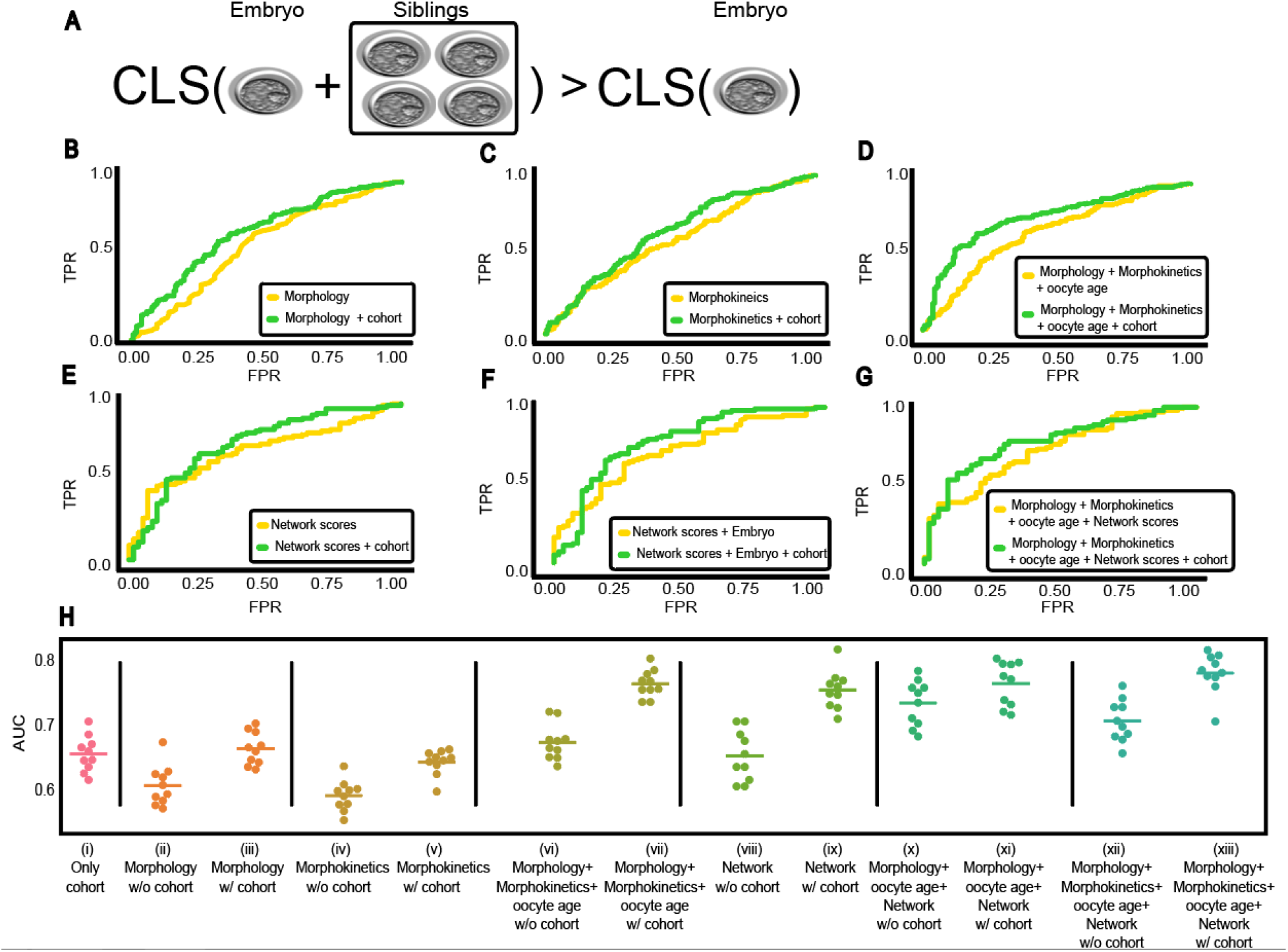
Implantation prediction performance (AUC) comparison of different model pairs with versus without sibling features establish that cohort properties contribute to implantation prediction. Statistical evaluation was computed with Wilcoxon signed rank test rejecting the null hypothesis that both models’ predictions are drawn from the same distribution. **(A)** Illustration: do cohort siblings data contribute to implantation prediction? **(B)** Morphology (Gardner scores). N=1936 embryos. N positive embryos = 1141, N negative embryos = 795. AUC: 0.6 versus 0.68, respectively, p-value < 0.0001. **(C)** Morphokinetics. N=2089 embryos. N positive embryos = 1176, N negative embryos = 913. AUC: 0.591 and 0.641, respectively, p-value < 0.0001. **(D)** Morphokinetics, morphology and oocyte age: N=1936 embryos. N positive embryos = 1141, N negative embryos = 795. AUC: 0.662 versus 0.764, respectively, p-value < 0.0001. **(E-G)** Deep convolutional neural network without (E) and with (F) morphology and oocyte age, and with morphokinetics, morphology and oocyte age (G). N=772 embryos. N positive embryos = 482, N negative embryos = 290 (**E**) AUC: 0.698 versus 0.744, respectively, p-value < 0.001. **(F)** AUC: 0.72 versus 0.779, respectively, p-value < 0.001. **(G)** AUC: 0.727 versus 0.8, respectively, p-value < 0.001. (**H**) Replication analysis. Performance assessment for models trained without versus with cohort features in ten independent partitioning to train and test sets. (i) Cohort features. AUC mean (standard deviation): 0.65 (0.02) (ii-iii) Morphology without (ii) or with (iii) cohort features. AUC mean (standard deviation) was 0.59 (0.02) without cohort versus 0.65 (0.02) with cohorts, Wilcoxon signed rank test p-value < 0.01. (iv-v) Morphokinetics without (iv) or with (v) cohort features. ACU mean (standard deviation) was 0.57 (0.02) without cohort versus 0.62 (0.01) with cohort, Wilcoxon signed rank test p-value < 0.01. Surprisingly, the model trained with morphokinetic and cohort features performed slightly worse than the model trained with cohort features alone (i versus v) perhaps due to inclusion of morphological features less correlative with the outcome without feature selection. (vi-vii) Morphokinetics, morphology and oocyte age without (vi) or with (vii) cohort features. AUC mean (standard deviation) was 0.65 (0.02) without cohort versus 0.74 (0.02) with cohort, Wilcoxon signed rank test p-value < 0.01. (viii-ix) Deep convolutional neural network scores without (viii) or with (ix) cohort features. AUC mean (standard deviation) 0.63 (0.03) without cohort versus 0.73 (0.02) with cohort, Wilcoxon signed rank test p-value < 0.01. (x-xi) Deep convolutional neural network scores with morphology and oocyte age without (x) or with (xi) cohort features. AUC mean (standard deviation) was 0.71 (0.03) without cohort versus 0.75 (0.03) with cohort, Wilcoxon signed rank test p-value < 0.01. (xii-xiii) Deep convolutional neural network scores with morphokinetics, morphology and oocyte age without (xii) or with (xiii) cohort features. AUC mean (standard deviation) was 0.69 (0.03) without cohort versus 0.77 (0.02) with cohort, Wilcoxon signed rank test p-value < 0.01.

Next, we turned to evaluate a deep convolutional neural model that extracts information directly from the raw embryo images. These “deep learning” models were shown to surpass more traditional machine learning models in many domains, including IVF embryo implantation prediction [13,14,15,16]. Specifically, we used a pre-trained VGG16 network [36] and fine-tuned it using preprocessed images of transferred blastocysts. Here too, we trained one model without cohort features, and another with the network’s confidence score along with cohort features (Methods). Similar to the previous models, inclusion of cohort features enhanced the model’s capacity to accurately predict implantation outcome (Fig. 3E). Moreover, cohort information enhanced the capacity to accurately predict implantation for a model that combined the deep learning model score, morphology features and the oocyte age (Fig. 3F), as well as for a model that also included morphokinetic features (Fig. 3G). Finally, we validated that these results were consistent by performing 10-fold cross validation: 10 rounds of training and evaluation for each model, each time with an independent partitioning of cohorts to train and test sets (Fig. 3H).These results, consistent across five models and multiple replicates, established that the siblings of the cohort encapsulate valuable information regarding the implantation potential of the transferred embryo.

### Identifying cohort properties driving the model’s prediction

While cohort information was found to enhance implantation prediction, it was not clear which of the cohort features contributed to the measured boost in performance. Thus, we next aimed toward explaining the models’ decisions. We focused our efforts on the two top performing models trained without or with the deep learning network’s feature (i.e., confidence score), namely, a model trained on morphology, morphokinetics and oocyte age. This enabled us, beyond plain model interpretability, to assess what information was encapsulated in the network beyond the morphology, morphokinetics and oocyte age.

To pinpoint what features were the most important for the model’s prediction we applied SHapley Additive exPlanations (SHAP), a game theory-based method for interpreting models’ predictions that assigns each feature an importance value for a prediction [37]. For the model that was trained solely with cohort features we found that the fraction of siblings reaching blastulation and the cohort size were the two most important features for implantation prediction (Fig. 4A). The trophectoderm score and expansion degree (morphology), oocyte age and seven morphokinetic features were identified as the ten top features in a model trained with morphokinetics, morphology and oocyte age (Fig. 4B). When comparing the ten top ranked features with a model trained with cohort features, we noticed that the fraction of sibling embryos reaching blastulation came up as the most important feature for prediction of implantation outcome, preceding morphology, morphokinetic and oocyte age (Fig. 4C). Overall, five of the top ten important features were attributed to the cohort and the other five were attributed to the transferred blastocyst (Fig. 4C). The network confidence score was the most informative feature, by a margin, in a model trained with the network confidence score, morphology (Gardner scores), morphokinetics and the oocyte age (Fig. 4D). Trophectoderm score was ranked only third and the expansion degree was not ranked within the ten most important features indicating that the network encoded the annotated morphology (Gardner scores). Cohort features were ranked higher than the morphology and morphokinetic features in a model that was trained with the network prediction, morphology, morphokinetics and the oocyte age (Fig. 4E). Half of the top features (5/10) were cohort-related suggesting that the cohort encodes information that is not included within the embryo’s image (Fig. 4E). While oocyte age was identified as an important feature in the two models that did not include cohort features (Fig. 4B and Fig. 4D), it was not one of the top ten features when cohort features were included (Fig. 4C and Fig. 4E). This suggests that the cohort encodes information more discriminative than the oocyte age in terms of implantation prediction. Altogether, these analyses show that cohort features are an important source of information for the prediction of implantation outcome.

**Figure 4:**
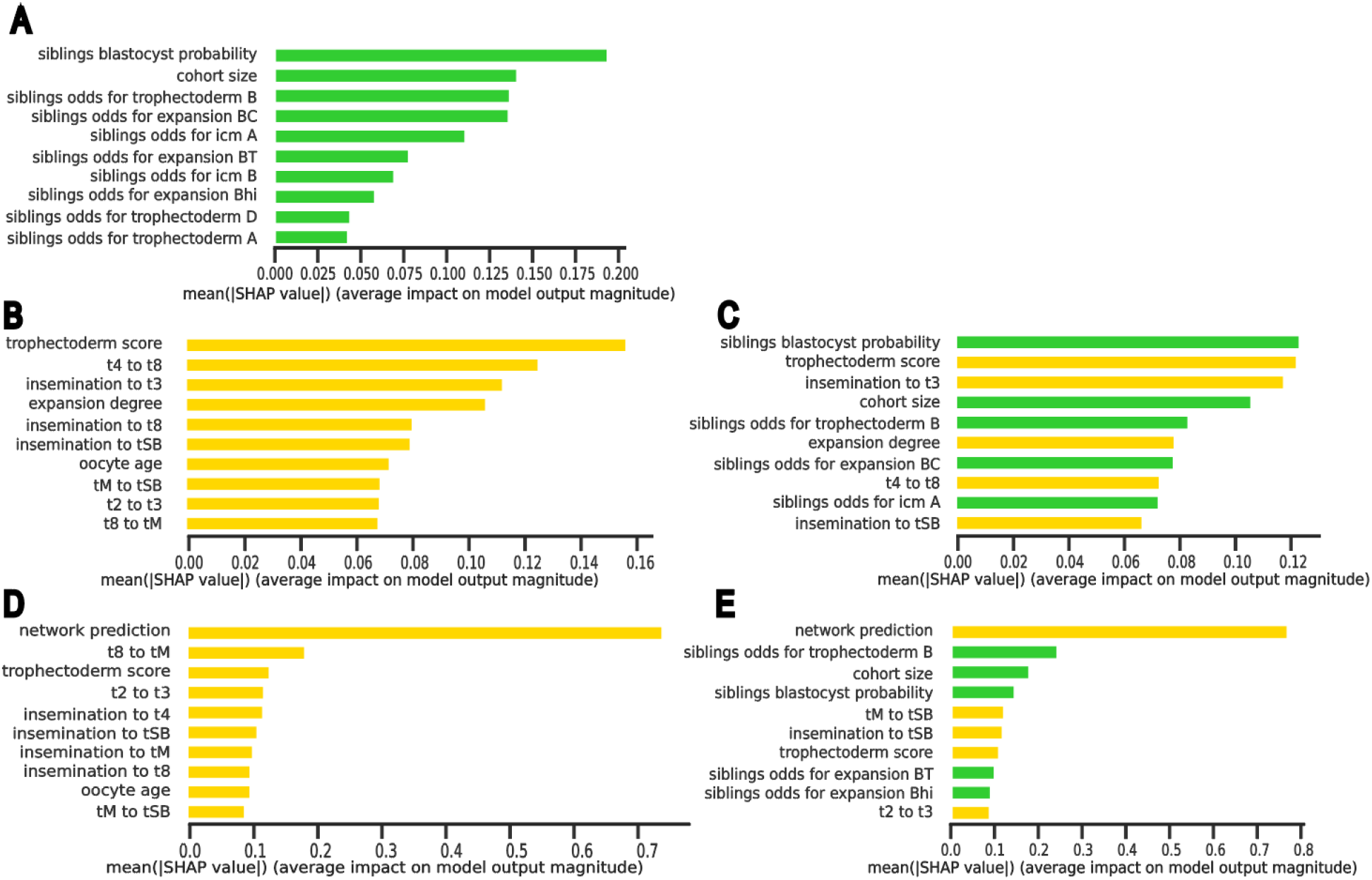
Model explainability analysis. Each panel shows features importance for the top ten features for a specific model using Shapely Additive Explanations (ShAP). (**A)** Only cohort features. (**B-C**) Morphokinetics, morphology and oocyte age without (**B**) or with (C) cohort features. (**D-E**) Deep convolutional neural network predictions with morphokinetics, morphology, oocyte age and without (D) or with (E) cohort features.

### Identifying cohort properties that corrected erroneous prediction

Finally, we evaluated the contribution of the cohort features to the classification of each of the transferred embryos. Fig. 5A shows the embryo classification scores by a model trained with morphology, morphokinetics and oocyte age without (x-axis) and with (y-axis) cohort features. Each data point corresponds to an embryo and the color code indicates positive (green) and negative (red) embryos. Embryos above the y = x diagonal had higher classification scores when including the cohort features, reflecting a higher prediction for successful implantation. Inclusion of cohort features increased the classification scores of positive embryos (green data points above the y = x diagonal) and decreased the classification scores of negative embryos (red data points below the y = x diagonal), indicating that adding cohort features to the model improves model’s discrimination for both positive and negative embryos (Fig. 5B). To better understand how cohort features enhanced implantation prediction we zoomed in to the embryos that were “rescued” by the cohort features, i.e., correctly classified only by the classifier that had access to cohort information. These included 15 negative and 62 positive “rescued” embryos. We examined the two top ranked cohort features, fraction of sibling embryos reaching blastulation and cohort size. We compared these features in cohorts of “rescued” embryos to all cohorts with the label corresponding to the “erroneous” prediction (by the model that did not have access to the cohort). For example, the two aforementioned cohort features of a positive embryo that was predicted as “negative” by a model that did not have access to cohort information and “rescued” by the cohort features, were compared to the corresponding features of all negative cohorts. This analysis should highlight whether these cohort features were correlated with the “rescued” prediction, and thus provide insight on what cohort features are used to improve the model’s prediction. While not identifying an obvious pattern for rescued negative embryos (Fig. 5C-D), we revealed in rescued positive embryos an elevation (relative to negative embryos) in the fraction of sibling embryos reaching blastulation and in the cohort size (Fig. 5E-F). A similar pattern, although less reliable due to the lower number of rescued embryos (because of the smaller dataset), was observed for models that included the neural network’s predictions (Fig. S2). These results suggest that these two cohort features were used by the model to correct negative-to-positive predictions but not positive-to-negative predictions.

**Figure 5:**
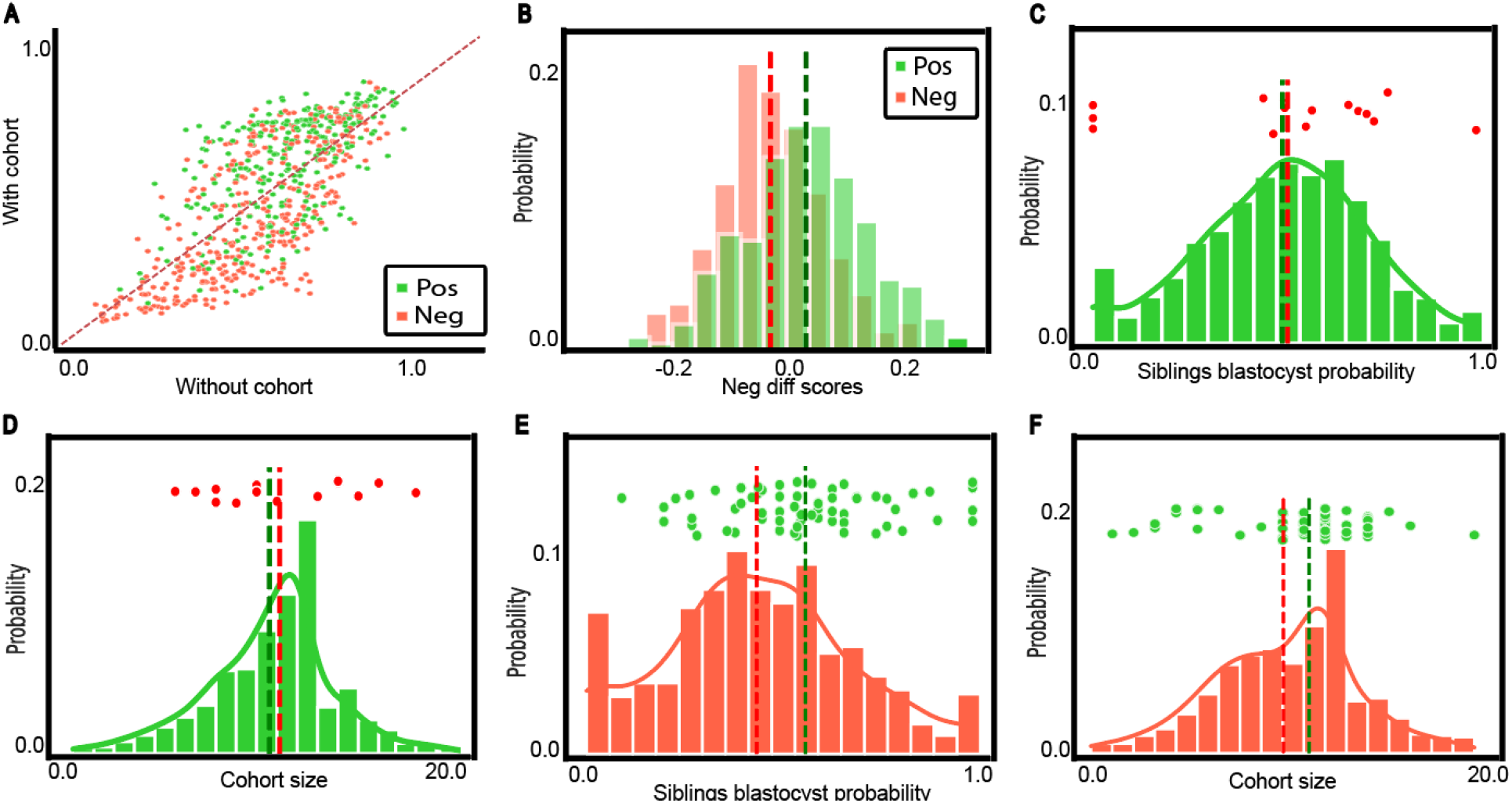
Analysis of cohort properties that “rescued” erroneous prediction. N transferred blastocysts =2089 from which 1176 were positive and 913 were negative embryos. The results refer to a model trained with morphology, morphokinetics and oocyte age without and with cohort features. **(A)** Embryos matched classification scores by the two models, without (x-axis) and with (y-axis) cohort features. **(B)** Distribution of the difference in the embryos matched classification scores: with - without cohort features. Mean (standard deviation) difference for positive cohorts was 0.043 (0.1) (Wilcoxon signed rank test p-value < 0.0001) versus -0.02 (0.09) (Wilcoxon rank-sum test p-value < 0.01) for negative cohorts. **(C-F)** Distribution of fraction of blastocysts siblings (C,E) or cohort size (D,F) for positive (green, C-D) or negative (red, E-F) embryos. Each of the data points above the distribution indicate an embryo that was “rescued” with the cohort feature, i.e., classified erroneously by a model trained without and corrected with a model trained with cohort features. **(C-D)** Negative embryos that were erroneously classified as positive without cohort features and were correctly classified by a model that had access to cohort features. N = 15 rescued embryos.(**C**) Mean (standard deviation) fraction of sibling embryos within a cohort (not including the transferred embryo/s) reaching blastulation was 0.49 (0.22) for positive cohorts versus 0.5 (0.29) for negative rescued embryos, Wilcoxon signed rank test on the differences from the positive embryos’ mean was not statistically significant. (**D**) Mean (standard deviation) cohort size was 10.56 (3.19) for positive cohorts versus 11.3 (3.64) for negative rescued embryos, Wilcoxon signed rank test on the differences from the positive embryos’ mean was not statistically significant. **(E-F)** Positive embryos that were erroneously classified as negative without cohort features and were correctly classified by a model that had access to cohort features. N = 62 rescued embryos. Distribution of the fraction of embryos within a cohort (not including the transferred embryo/s) to develop to a blastocyst (E) or cohort size (i.e., number of sibling embryos in a cohort) (F) compared across negative embryos versus positive embryos that were “rescued” by the cohort features, i.e., correctly classified only by the classifier that had access to cohort information. (**E**) Mean (standard deviation) fraction of sibling embryos within a cohort (not including the transferred embryo/s) reaching blastulation was 0.41 (0.25) for negative cohorts versus 0.56 (0.21) for positive rescued embryos, Wilcoxon signed rank test on the differences from the negative embryos’ mean p-value < 0.0001. (**F**) Mean (standard deviation) cohort size was 9.9 (3.36) for negative cohorts versus 10.74 (3.17) for positive rescued embryos, Wilcoxon signed rank test on the differences from the negative embryos’ mean p-value < 0.01.

## Discussion

In vitro fertilization (IVF) is a perfect system for studying the effect of genotypic and environmental variation on phenotype. This is due to the availability of high-content human embryo data that includes phenotypic information regarding multiple sibling embryos for each treatment cycle, who share a common genetic background and similar external conditions. From the machine learning perspective, IVF is a fitting example for an application where vast unlabeled data, specifically from non-transferred cohort siblings, can provide valuable information for a more accurate prediction of embryo phenotypic quality, i.e., implantation potential. These biological and machine learning concepts converge to a common theme where the uncertainty in the transferred embryo features, due to either inconsistency in annotations, features that were not explicitly measured or label ambiguity, can be reduced by information encapsulated in the correlated cohort embryos. We believe that this is achieved by noise reduction with the multiple correlated instances.

We established that embryos from the same cohort were more phenotypically similar than embryos from different cohorts (Fig. 1), demonstrated that siblings of successfully implanted (positive) embryos were of higher phenotypic quality in relation to siblings of negative embryos (Fig. 2), and demonstrated that cohort features contribute to machine learning based implantation prediction (Fig. 3). The latter was achieved by extracting a new set of features from unlabeled siblings within the cohort, incorporating them with different feature sets, and comparing the classifier’s performance in implantation prediction without versus with cohort features. Even though each individual cohort feature had only a marginal effect (Fig. 2), the machine learning driven integration of all cohort features led to a consistent improvement in implantation prediction regardless of the embryo-focused model (Fig. 3). These results suggest a general concept where the transferred embryo’s siblings encapsulate discriminative information that is complementary to the information encoded in the transferred embryo, and thus, cohort features is likely to contribute to any embryo-derived features. Since the siblings’ data are routinely collected in the clinic, incorporating cohort features in AI-driven embryo implantation prediction can have direct translational implications in the clinic.

Previous studies correlated cohort-based properties to implantation outcome. For example, demonstrating improved outcome for embryos selected from cohorts with more than five embryos [25], or from day 3 cohorts where at least one sibling embryo achieved blastulation after extended culture [28,29]. Other studies incorporated specifically designed cohort-based features to machine learning models, specifically the cohort size [24,27] and number of developed embryos [33], or even incorporated multiple cohort-related features to show that the cohort alone contains discriminative information [26]. We performed a comprehensive analysis that systematically assessed the contribution of incorporation of cohort properties to existing models for prediction of implantation outcome. We did not decide on specific thresholds, rather we provided all the “raw” features to the AI machinery to automatically determine and weight which and what combination was most discriminative. This unbiased approach allowed us to reveal how discriminative each cohort property is in the context of a given model (Fig. 4) and which cohort properties were critical to “rescue” embryos that were incorrectly classified without cohort properties (Fig. 5). Importantly, most of the previous studies mentioned above assessed embryos that were transferred at day 3 from fertilization. At this early stage of embryo development, the uncertainty of the implantation potential of a single embryo is higher [3839], and thus, additional correlated measurements in cohort features are expected to provide a more discriminative signal. This is especially relevant when sibling embryos are kept in extended blastocyst culture and their blastulation outcomes are known. Our results relate to blastocysts, i.e., day 5 embryos, when the uncertainty is lower. Still, we were able to establish that the cohort information contributes discriminative power beyond the transferred embryo features.

We found that cohort features “rescued” 4-fold more positive (N = 62) versus negative (N = 15) embryos (Fig. 5). One explanation for this asymmetry could be due to the ambiguity in the negative labels. While positive embryos are inherently of high implantation quality, negative embryos fail implantation because of either low quality embryos and/or poor endometrial or uterine factors. This creates uncertainty in the ground truth labels of non-implanted embryos, or “label ambiguity” [20]. Thus, it could be easier to “rescue” a positive embryo that was mistakenly classified as “negative” with a quality cohort, in comparison to a negative embryo that could be of high implantation potential (also encoded in its cohort features).

In this study, digital embryo images were manually annotated for morphology (based on the Gardner embryo scoring system) and key morphokinetic events using the time-lapse information. Manual annotation is throughput-limiting. Automated tools for morphological evaluation [22,23] and detection of morphokinetic events [2040] are quickly reaching human-level performance and have the advantage of avoiding intra- and inter-annotator bias potentially replacing the need for manual annotations in the near future.

In the context of machine learning, this is an example of semi-supervised learning, where a small fraction of labeled (transferred embryos) and vast unlabeled observations (non-transferred cohort siblings) are used together to improve the learned model’s performance toward the task of predicting implantation potential. While the underlying assumption in semi-supervised learning is that the observations are unrelated, here we characterize another type of semi-supervised learning, where the unlabeled observations are associated with the labeled observations, and suggest a way to use information from these unlabeled observations for improving the prediction power of the supervised models.

## Methods

### Experiments

#### Data collection and ethics

The data included for this analysis were retrospectively collected from IVF cycles conducted at a single center between March 2010 and December 2018. Historical images of blastocyst-stage embryos and metadata were provided by AiVF. All procedures and protocols were approved by an Institutional Review Board for secondary research use (IRB reference number HMO-006-20). All IVF cycles were with either the patient’s own oocytes (n=1134) or with donor oocytes (n=955), ages 18-51 years old (Fig. S3). In cases of oocyte donation, donor age was considered as the “oocyte age”.

Fertilization was determined by the presence of two pronuclei (2PN) 16-18 hours after insemination. All zygotes were placed inside the EmbryoScope™ time-lapse incubator system (Vitrolife, Denmark) and incubated using sequential media protocol until blastocyst-stage. Image acquisition using the EmbryoScope™ imaging software occurred every 15-20 minutes. For every embryo, seven-layers of z-stack images 15µm apart were acquired at each time point, where time 0 was defined as the fertilization time. A total of 16,194 oocytes from 1,605 IVF cycles were recorded. All cycles included at least one fresh single embryo transfer at blastocyst stage with a known implantation outcome (implanted/not implanted). Out of 2089 embryos with known implantation outcomes, 1176 were successfully implanted (*positive embryo*s) and 913 failed to implant (*negative embryos*).

#### Annotation of embryo clinical quality

Following in vitro incubation, the number of embryos from the cohort transferred was either one or two. In cycles where two embryos were transferred, we only included cases where both embryos either successfully implanted or both failed implantation. Implantation following embryo transfer was determined by ultrasound scanning for gestational sac after ∼seven weeks of pregnancy. Positive embryos (and their corresponding positive cohorts) were defined when the number of gestational sacs matched the number of transferred embryos. Negative embryos (and their corresponding negative cohorts) were defined when no gestational sac following embryo transfer was observed.

### Analysis

#### Embryo morphological feature representation

Embryos were morphologically annotated at the blastocyst stage (day 5 of embryonic development) according to the Gardner scoring scheme by onsite trained embryologist ∼120 hours post insemination[32,8]. The time of blastulation was determined by time-lapse monitoring of embryo development. Specifically, every embryo was assigned a three-part alphanumeric quality score based on its expansion status (“blastocyst expansion”, ranked 1-6), morphology of the inner cell mass (“ICM”, ranked A-D), and morphology of the trophectoderm (“TE”, ranked A-D) (Fig. 1H). Embryos with missing Gardner annotations were excluded from the analysis.Out of the 2089 embryos, 1936 had Gardner annotations, where 1141 were annotated as positive and 795 as negative. Embryos’ morphological variables were computationally represented via one-hot encoding, i.e., a feature vector of size 6 + 4 + 4 = 14 representing the Gardner scores.

#### Embryo morphokinetic feature representation

Time-lapse images of embryo development were viewed by onsite trained embryologist and seven key morphokinetic events were manually annotated in accordance with published consensus criteria[42]: Time of fertilization (t0), cell division to the 2,3,4, and 8-cell stage (t2, t3, t4, t8), compaction of the morula (tM), and time of blastulation (tSB). Missing annotations due to limitations of the dataset were completed according to the following rules, as determined by domain experts: t2 = time of pronuclei disappearance (tPNf - time when both pronuclei disappear, independently annotated, see below) + 2 hours; t3 = t4 - 1 hour; t4 = t3 + 1 hour; t8 = t7 + 3 hours; tM = tSB - 6 hours. Missing morphokinetic event annotations were at levels of 5%- 10%, except for tM and tSB with 20% missing annotations. After manually completing the missing annotations, embryos with a further single missing value were determined by first finding the five most similar embryos based on the remaining available annotated morphokinetic features and then using their mean value of the missing features. Overall, the morphokinetic feature vector was of size 11 and included the (five) time intervals between every consecutive developmental stage and the (six) overall time points for each developmental stage from the time of fertilization. Importantly, our dataset also included manual annotations beyond the morphokinetic events listed above (e.g., tPNf, see above). These events were not included in the morphokinetic feature representation used in this analysis because many embryos in the dataset did not have these morphokinetic events annotated.

#### Similarity between timing of morphokinetic events

We compared the similarity between the timings of morphokinetic events and between the timings of consecutive morphokinetic events between embryo pairs within the same cohort (siblings) and in different cohorts (non siblings). For each morphokinetic event timing and consecutive morphokinetic events timing, we compared the similarity of all sibling and non-sibling pairs. A similarity measure that encodes the full morphokinetic profile was calculated as the Euclidean distance between normalized vectors that included the timing of all the morphokinetic events and the timing of all the consecutive morphokinetic events.

To compare morphokinetic similarities between sibling and non sibling embryo pairs with similar morphological properties, we evaluated all embryo triplets that share the same Gardner scores, where two embryos were siblings and the third embryo was from a different cohort.For each triplet we evaluated the similarity in morphokinetics between the siblings versus the non siblings pairs.

#### Cohort morphology-based feature representation

A cohort contains multiple embryos from the same couples in the same IVF treatment cycle. Seventeen (17) cohort features were extracted from all cohort siblings, excluding the transferred embryo. These included the cohort size (the only feature that included the transferred embryo, the fraction of sibling embryos reaching blastulation (Fig. 2B), the fraction of sibling embryos that hatched, a 13-dimensional vector encoding the fraction of siblings for each Gardner score (5+4+4 features).

#### Automated deep learning based embryo implantation prediction

We had access to historical images of 772 transferred blastocyst-stage embryos from 638 cohorts and associated known implantation outcomes. Of these, 482 embryos successfully implanted (positive) and 290 failed to implant (negative).

##### Images Preprocessing

Our analysis focused on the blastocyst’s last time-frame prior to hatching and the center z-stack image.

We preprocessed the raw frames in order to segment the embryo region from the surrounding well and background. This allowed us to train more complex models on a large training set in reasonable time while keeping the inherent spatial resolution features of the embryo. To localize and segment the embryo we developed the following pipeline. First, a mask-RCNN [43] was trained to identify a bounding box around each embryo using 800 manually annotated images.Second, a hough-transform [44] was applied to center the embryo in the image by detecting a circular object within the bounding box. Third, a U-NET model was trained based on 500 manually validated outputs from the previous step to provide the embryo segmentation mask [45]. Our U-NET architecture consisted of 4 convolutional layers for the encoder (downsampling) / decoder (upsampling) with 32, 64, 128 and 256 filters correspondingly. Each layer included batch normalization and relu activation function with maxpooling for the encoder and upsampling for the decoder. Finally, the image was further resized to 64×64. The trained pipeline architecture is presented in Fig. S4.

##### Image classification model

A pretrained VGG16 [36] architecture was used as a backbone followed by a flattening layer, a fully connected 16 node dense layer and a dense single node layer with a sigmoid activation. The full model was retrained (no freezing of weights) using binary cross entropy loss and the Adam optimizer with a learning rate of 0.00001 to output a confidence score in the range of 0-1 for predicting implantation probability.

The model was trained for 100 epochs with a batch size of 32. For each batch, image-augmentation (brightness, flipping, rotation and Gaussian noise) was performed prior to fitting the model to reduce overfit. We trained the model with 10-fold cross validation allowing all the available data points to run at inference and output a confidence score.

#### Evaluation of machine learning models without versus with cohort features

Models were trained with morphology features and/or morphokinetic features and/or the oocyte age. Each feature representation was used to train and then evaluate the performance of two models: without including cohort features (i.e., using only the transferred embryo information) versus including cohort features. The deep learning model was used as a feature extractor, where the model confidence score was one feature, and this features were used together with other features (e.g., morphokinetics, oocyte age) to train and then evaluate the performance of models that had vs. those did not have access to cohort features. Models based on the same features without or with cohort features were compared to evaluate the contribution of cohort features on the overall prediction. The data was partitioned to train (80%) and test (20%) sets maintaining a similar ratio of positive (i.e., successful implantation) and negative (failed implantation) cohorts in both sets. An XGBoost binary classification model was trained for each feature set [46]. When included, the confidence score of the deep learning model was considered as a single feature.Receiver Operating Characteristic (ROC) curve (i.e., true positive rate (TPR, sensitivity) versus false positive rate (FPR, 1-specificity)) and its Area Under the Curve (AUC) were used to visually and quantitatively compare performance of model pairs without versus with cohort features. Hyper-parameter tuning was performed independently for each XGBoost classifier using sklearn’s GridSearchCV [47]. The parameters that were optimized were: (1) Number of gradient boosted trees, (2) Minimum loss reduction required to make a further partition on a leaf node of the tree, (3) Maximum tree depth for base learners, (4) Subsample ratio of the training instance, (5) Subsample ratio of columns when constructing each tree, and (6) Minimum sum of instance weight (hessian) needed in a child.

Due to the lower volume of annotated raw image data, we applied feature selection, independently for each model, with the extremely Randomized Trees Classifier (ExtraTrees) classifier (an updated version of random forest) [48] to reduce the number of features to ten (unless the number of features was already lower).

#### Feature importance analysis

For interpretability purposes, we applied the feature importance method with Shapley Additive Explanations (SHAP) [**Error! Reference source not found**.37]. SHAP computes the contribution of each feature to each individual prediction. Embryo morphological categorical parameters were transformed into ordinal variables for the SHAP analysis.

## Funding and Acknowledgments

This research was supported by the Israel Council for Higher Education (CHE) via the Data Science Research Center, Ben-Gurion University of the Negev, Israel. We thank Nadav Rappoport, Zeev Bomzon, and Yishay Tauber for critically reading the manuscript.

## Author Contribution

AZ conceived the study. NT and OR developed analytic tools, and analyzed the data. NT, OR and AZ interpreted the data and drafted the manuscript. MTS, RM, MM, DG and DSS provided clinical input for the presentation of the manuscript, reviewed and revised the manuscript. AZ mentored NT and OR. All authors edited the manuscript and approved its content.

## Competing Financial Interests

OR, MTS, RM, DG, and DSS are employees at AIVF LTD.

## Supplementary Tables

**Table S1:** Number of transferred blastocysts, number of implanted blastocysts (positive embryos) and non-implanted blastocysts (negative embryos), number of cohorts, number of positive and negative cohorts and number of cohort’s siblings for each dataset.

|  | # embryos | # implanted embryos | # non-implanted embryos | # cohorts | # positive cohorts | # negative cohorts | # siblings |
| --- | --- | --- | --- | --- | --- | --- | --- |
| **Morphokinetics annotated** | 2089 | 1176 | 913 | 1605 | 928 | 677 | 14105 |
| **Morphology annotated** | 1936 | 1141 | 795 | 1513 | 908 | 605 | 13493 |
| **Image data** | 772 | 482 | 290 | 638 | 404 | 234 | 5990 |

## Supplementary Figures

**Figure S1.**
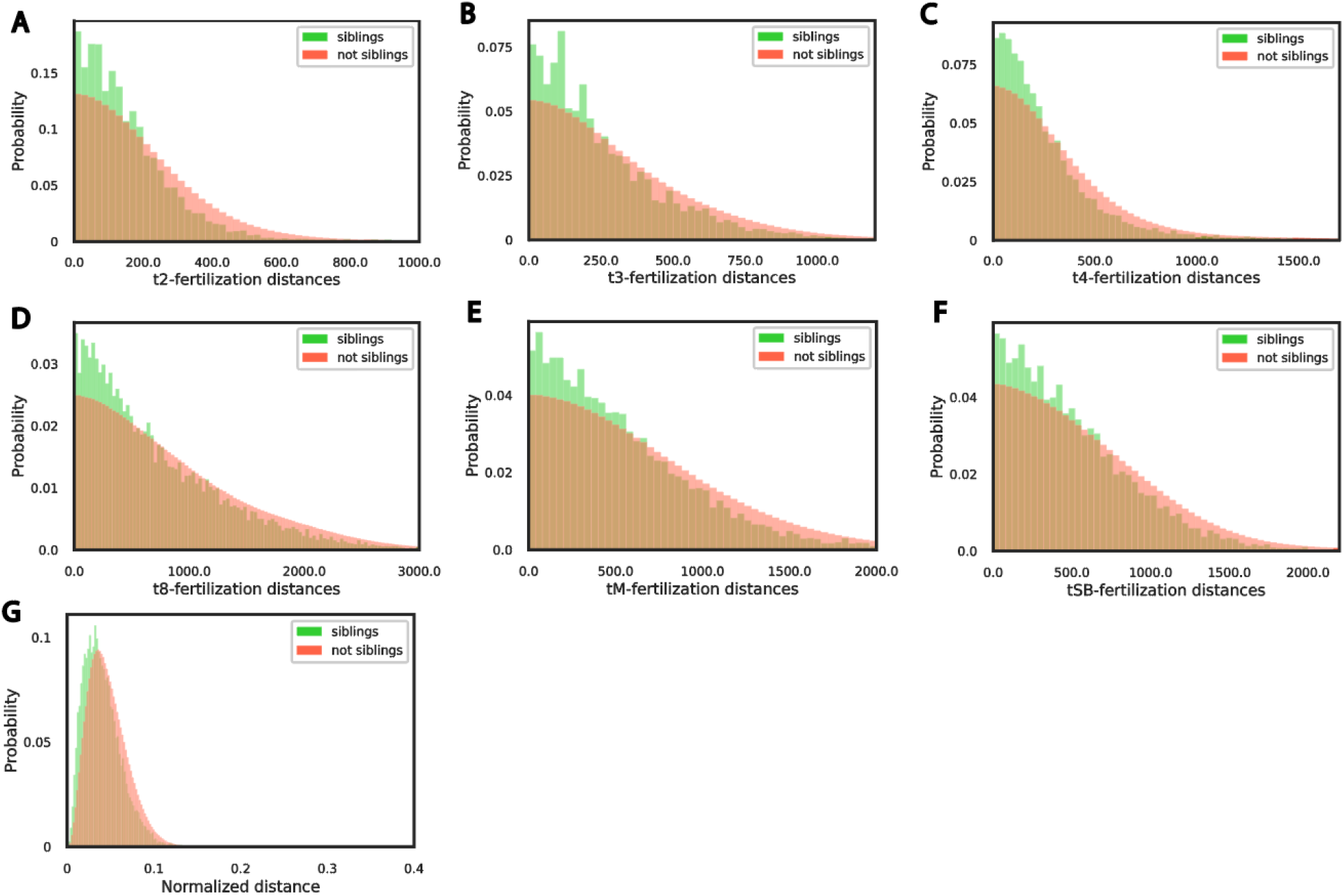
Sibling embryos from the same cohort are more similar than non-siblings. (**A-F**) Distribution of the difference in time intervals, in minutes (A-F) or normalized distance (G), between fertilization and each morphokinetic event compared across siblings versus not siblings embryo pairs. morphokinetic features: cell division to the 2, 3, 4 and 8-cell stage (t2, t3, t4, t8), the compaction of the morula - a day-3 development stage (tM) and the start of blastulation (tSB) - a day-5 development stage. N embryos = 16194. N cohorts = 1605. N positive cohorts = 928, N negative cohorts = 677. (**A**) Mean (standard deviation) of distances between fertilization-t2 intervals was 158.56 (150.59) for sibling embryos versus 217.62 (218.51) for not-sibling embryos, Mann-Whitney-U signed rank test p-value < 0.0001. (**B**) Mean (standard deviation) of distances between fertilization-t3 intervals was 258.21 (238.36) for sibling embryos versus 327.68 (290.56) for not-sibling embryos, Mann-Whitney-U signed rank test p-value < 0.0001. (**C**) Mean (standard deviation) of distances between fertilization-t4 intervals was 261.45 (268.56) for sibling embryos versus 333.01 (314.93) for not-sibling embryos, Mann-Whitney-U signed rank test p-value < 0.0001. (**D**) Mean (standard deviation) of distances between fertilization-t8 intervals was 700.46 (595.04) for sibling embryos versus 836.65 (666.99) for not-sibling embryos, Mann-Whitney-U signed rank test p-value < 0.0001. (**E**) Mean (standard deviation) of distances between fertilization-tM intervals was 537.38 (443.54) for sibling embryos versus 655.45 (515.67) for not-sibling embryos, Mann-Whitney-U signed rank test p-value < 0.0001. (**F**) Mean (standard deviation) of distances between fertilization-tSB intervals was 499.29 (397.18) for sibling embryos versus 596.34 (462.68) for not-sibling embryos, Mann-Whitney-U signed rank test p-value < 0.0001. (**G**) Mean (standard deviation) of normalized distances between all morphokinetic features time intervals from fertilization was 0.04 (0.02) for sibling embryos versus 0.05 (0.02) for not-sibling embryos, Mann-Whitney-U signed rank test p-value < 0.0001.

**Figure S2:**
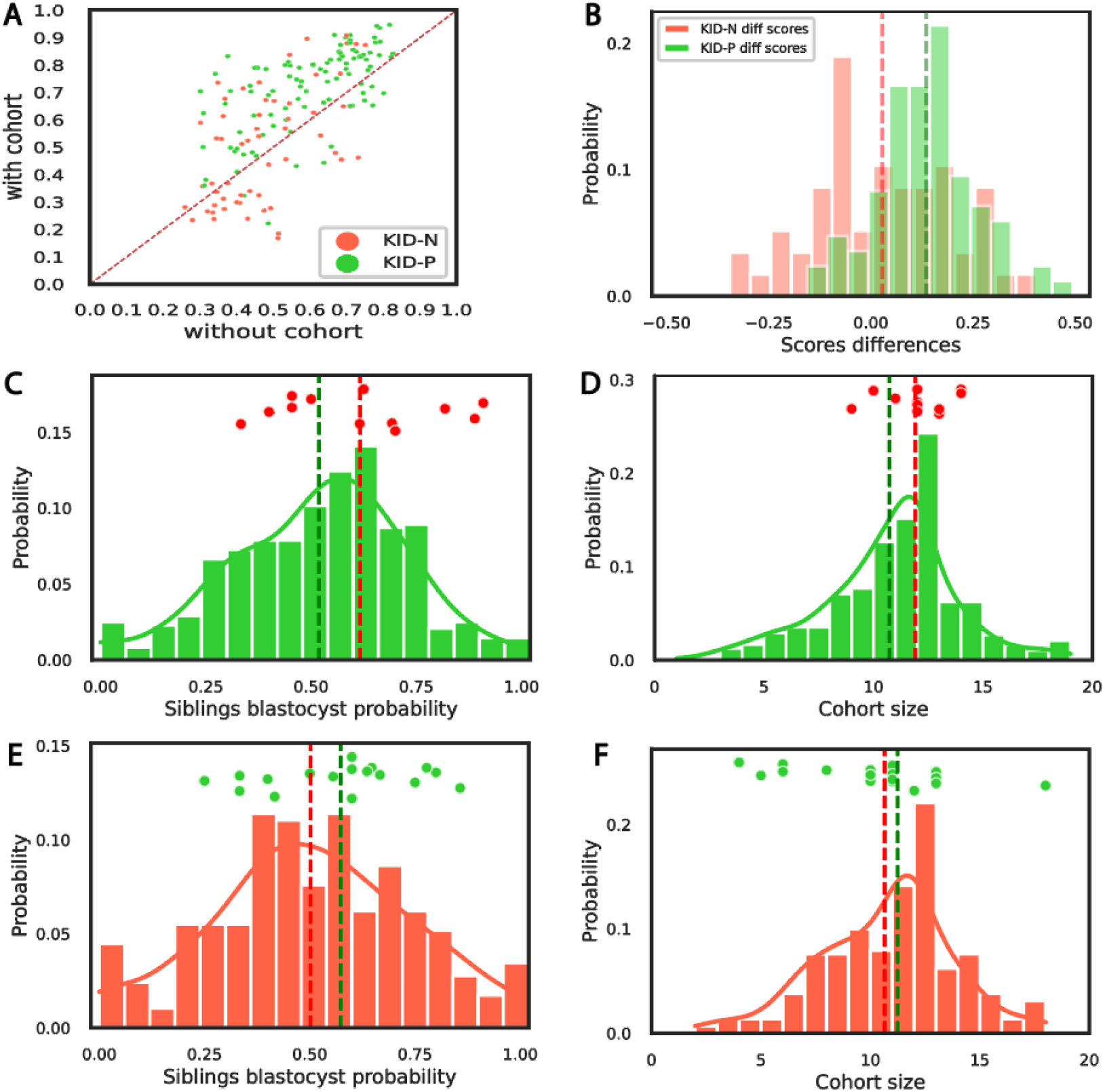
Analysis of cohort properties that “rescued” erroneous prediction. N transferred blastocysts = 772 from which 482 were positive and 290 were negative embryos. The results refer to a model trained with deep convolutional neural network, morphology, morphokinetics and oocyte age without and with cohort features. **(A)** Embryos matched classification scores by the two models: without (x-axis) and with (y-axis) cohort features. **(B)** Distribution of the difference in the embryos matched classification scores: with - without cohort features. Mean (standard deviation) difference for positive cohorts was 0.13 (0.11) (Wilcoxon rank-sum test p-value < 0.0001) versus 0.01 (0.17) (Wilcoxon rank-sum test was not statistically significant) for negative cohorts. **(C-F)** Distribution of fraction of blastocysts siblings (C,E) or cohort size (D,F) for positive (green, C-D) or negative (red, E-F) embryos. Each of the data points above the distribution indicate an embryo that was “rescued” with the cohort feature, i.e., classified erroneously by a model trained without and corrected with a model trained with cohort features. **(C-D)** Negative embryos that were erroneously classified as positive without cohort features and were correctly classified by a model that had access to cohort features. N = 12 rescued embryos.(**C**) Mean (standard deviation) fraction of sibling embryos within a cohort (not including the transferred embryo/s) reaching blastulation was 0.51 (0.2) for positive cohorts versus 0.61 (0.19) for negative rescued embryos, Wilcoxon signed rank test on the differences from positive embryos mean found no statistical significance. (**D**) Mean (standard deviation) cohort size was 10.56 (3.19) for positive cohorts versus 11.91 (3.64) for negative rescued embryos, Wilcoxon signed rank test on the differences from positive embryos mean found no statistical significance. **(E-F)** Positive embryos that were erroneously classified as negative without cohort features and were correctly classified by a model that had access to cohort features. N = 18 rescued embryos. Distribution of the fraction of embryos within a cohort (not including the transferred embryo/s) to develop to a blastocyst (E) or cohort size (i.e., number of sibling embryos in a cohort) (F) compared across negative embryos versus positive embryos that were “rescued” by the cohort features, i.e., correctly classified only by the classifier that had access to cohort information. (**E**) Mean (standard deviation) fraction of sibling embryos within a cohort (not including the transferred embryo/s) reaching blastulation was 0.5 (0.23) for negative cohorts versus 0.57 (0.17) for positive rescued embryos, Wilcoxon signed rank test on the differences from negative embryos mean found no statistical significance (p-value = 0.07). (**F**) Mean (standard deviation) cohort size was 10.64 (3.03) for negative cohorts versus 10.11 (3.39) for positive rescued embryos, Wilcoxon signed rank test on the differences from negative embryos mean found no statistical significance.

**Figure S3:**
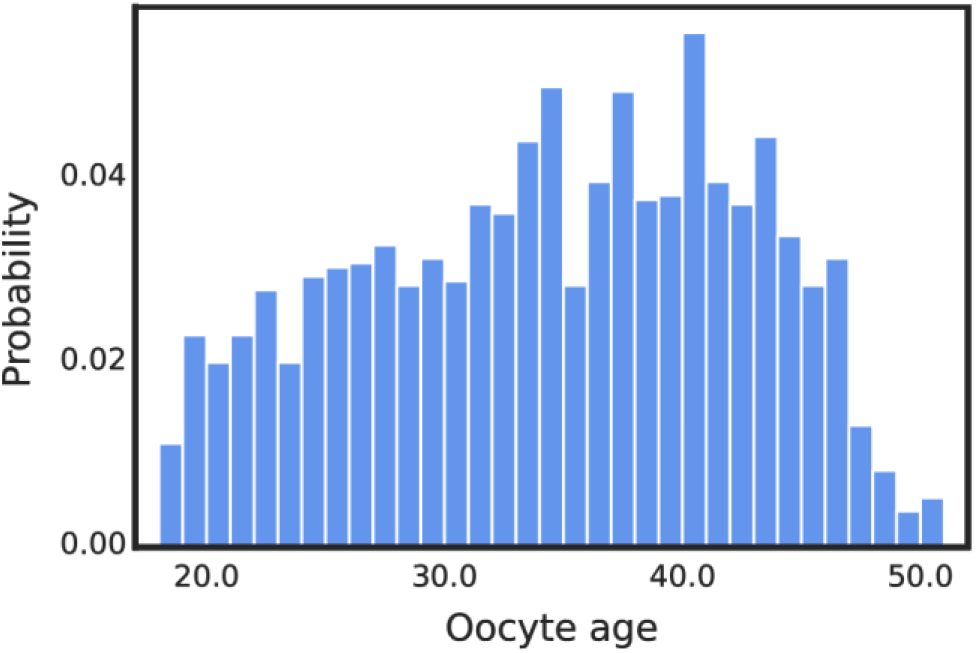
Distribution of oocyte age for all treatments. N= 2089. N cohorts = 1605. Mean (standard deviation) oocyte age was 33.99 (7.98).

**Figure S4:**
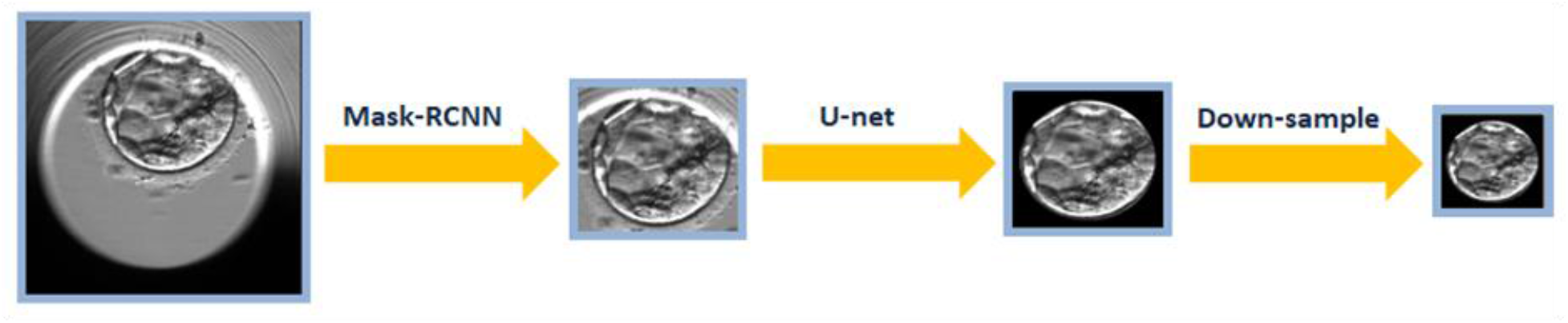
Oocyte segmentation pipeline (left-to-right). Mask-RCNN detects the embryo’s bounding box. The cropped image is segmented by a U-NET network. Finally the image is down sampled to 64×64.

